# DiCARN-DNase: Enhancing Cell-to-Cell Hi-C Resolution Using Dilated Cascading ResNet with Self-Attention and DNase-seq Chromatin Accessibility Data

**DOI:** 10.1101/2024.10.31.621380

**Authors:** Samuel Olowofila, Oluwatosin Oluwadare

## Abstract

The spatial organization of chromatin is fundamental to gene regulation and essential for proper cellular function. The Hi-C technique remains the leading method for unraveling 3D genome structures, but the limited availability of high-resolution Hi-C data poses significant challenges for comprehensive analysis. Deep learning models have been developed to predict high-resolution Hi-C data from low-resolution counterparts. Early CNN-based models improved resolution but struggled with issues like blurring and capturing fine details. In contrast, GAN-based methods encountered difficulties in maintaining diversity and generalization. Additionally, most existing algorithms perform poorly in cross-cell line generalization, where a model trained on one cell type is used to enhance high-resolution data in another cell type. In this work, we propose DiCARN (Dilated Cascading Residual Network) to overcome these challenges and improve Hi-C data resolution. DiCARN leverages dilated convolutions and cascading residuals to capture a broader context while preserving fine-grained genomic interactions. Additionally, we incorporate DNase-seq data into our model, providing a robust framework that demonstrates superior generalizability across cell lines in high-resolution Hi-C data reconstruction. DiCARN is publicly available at https://github.com/OluwadareLab/DiCARN

## 1 Introduction

Chromosome Conformation Capture (3C) technology is a molecular method used to analyze the spatial organization of chromatin in a cell (Lieberman-Aiden *et al*., 2009). The technology provides insights into the three-dimensional (3D) architectural arrangement of chromosomes, allowing researchers to study the physical interactions between DNA segments that may be separated by large genomic distances along the linear genome. In recent genetics research, high-throughput chromosome conformation capture (Hi-C) has emerged as the preferred 3C technique for deciphering and analyzing spatial genome organization within the eukaryotic cell nuclei. It is a genome-wide approach to the study of three-dimensional chromatin conformation inside the nucleus ((Lieberman-Aiden *et al*., 2009)). Lately, Hi-C has been the trailblazer technique in the exploration and characterization of genomic structural components, including A/B compartments, TADs (Dixon *et al*., 2012), frequently interacting regions (FIREs) (Schmitt *et al*., 2016), stripes (Vian *et al*., 2018), and enhancer-promoter interactions (Rao *et al*., 2014). Being a biochemical approach that allows for an all-versus-all mapping of chromosomal and genome fragment interactions, Hi-C takes into account the interaction between pair-read assays generated from a wet lab process, resulting in a symmetric (*n* x *n*) contact matrix representation of the Interaction Frequencies (IF), where *n* is the number of cells or evenly sized divisions of the genome called bins. The number indicated in every matrix cell represents the count of paired-end reads across two bins. The sizes of these bins, also known as ‘resolution,’ habitually range from 1 kiloBase (KB) to 2.5 MegaBase (MB), whose range hinges on the sequencing depth. The relevance of Hi-C data is spiking geometrically owing to its practicability in elucidating the genome organization. (Oluwadare *et al*., 2019)

However, a critical challenge in this research domain, is the limited availability of the required Hi-C resolution for exhaustive studies of genomic structures. This challenge has inspired the use of Deep Learning (DL) models to predict the required high-resolution Hi-C data from the more readily available low-resolution variants, sharing interest similarities with the Single-Image SuperResolution problem in the computer vision domain (He *et al*., 2016).

Zhang et al. (2018) pioneered high-resolution Hi-C data prediction with HiCPlus (Y. Zhang *et al*., 2018), a CNN-based model inspired by SRCNN (Dong *et al*., 2014), which used a three-layer CNN to impute high-resolution interaction frequencies. HiCNN (Liu and Z. Wang, 2019), a 54-layer CNN was modeled after DRRN (Tai *et al*., 2017). Both methods laid the groundwork for using CNNs in Hi-C enhancement, subsequent models focused on addressing challenges in improving resolution and generalization. SRHiC (Z. Li and Dai, 2020) introduced a ResNet-based approach (He *et al*., 2016) for Hi-C data enhancement, followed by the 2020 development of GAN-based models like DeepHiC (Hong *et al*., 2020) and HiCSR (Dimmick, 2020), which improved resolution enhancement by utilizing generator-discriminator networks. Later, HiCARN (Hicks and Oluwadare, 2022) introduced a more efficient cascading GAN, while DFHiC (B. Wang *et al*., 2023) advanced the field with a dilated full convolution network, preserving positional information and addressing previous shortcomings.

Despite these improvements, challenges such as limited receptive fields, lack of global context, and instabilities, particularly with mode collapse in GAN-based methods like HiCSR and DeepHiC, persist. Mode collapse results from the generator’s failure to produce diverse, representative samples, leading to incomplete data reconstruction. Additionally, most existing approaches have focused on architectural enhancements without integrating biologically relevant data, such as chromatin accessibility data, that reveals the chromosomal regions actively involved in gene regulation, which could provide more robust HR enhancement. In addition, existing algorithms perform poorly for cross-cell line generalization, where a model is trained on one cell and used for high-resolution enhancement of another cell. This limitation significantly affects the model’s scalability and applicability in broader biological research, where variability across cell lines is common.

In this work, we propose DiCARN (Dilated Cascading Residual Network), a novel approach to overcoming these challenges. DiCARN improves model stability by employing dilated convolutions for a larger receptive field and incorporates chromatin accessibility data, enabling more accurate and biologically meaningful Hi-C resolution enhancement.

## 2 Materials and Method

### 2.1 Architecture

Our proposed model, DiCARN, implements a novel fusion of dilated convolutions, spatial selfattention ((Vaswani *et al*., 2017; Q. Zhang *et al*., 2022)), and cascaded residual networks ((Ahn *et al*., 2018)), with its visual outlay depicted in Figure 1.

**Figure 1:**
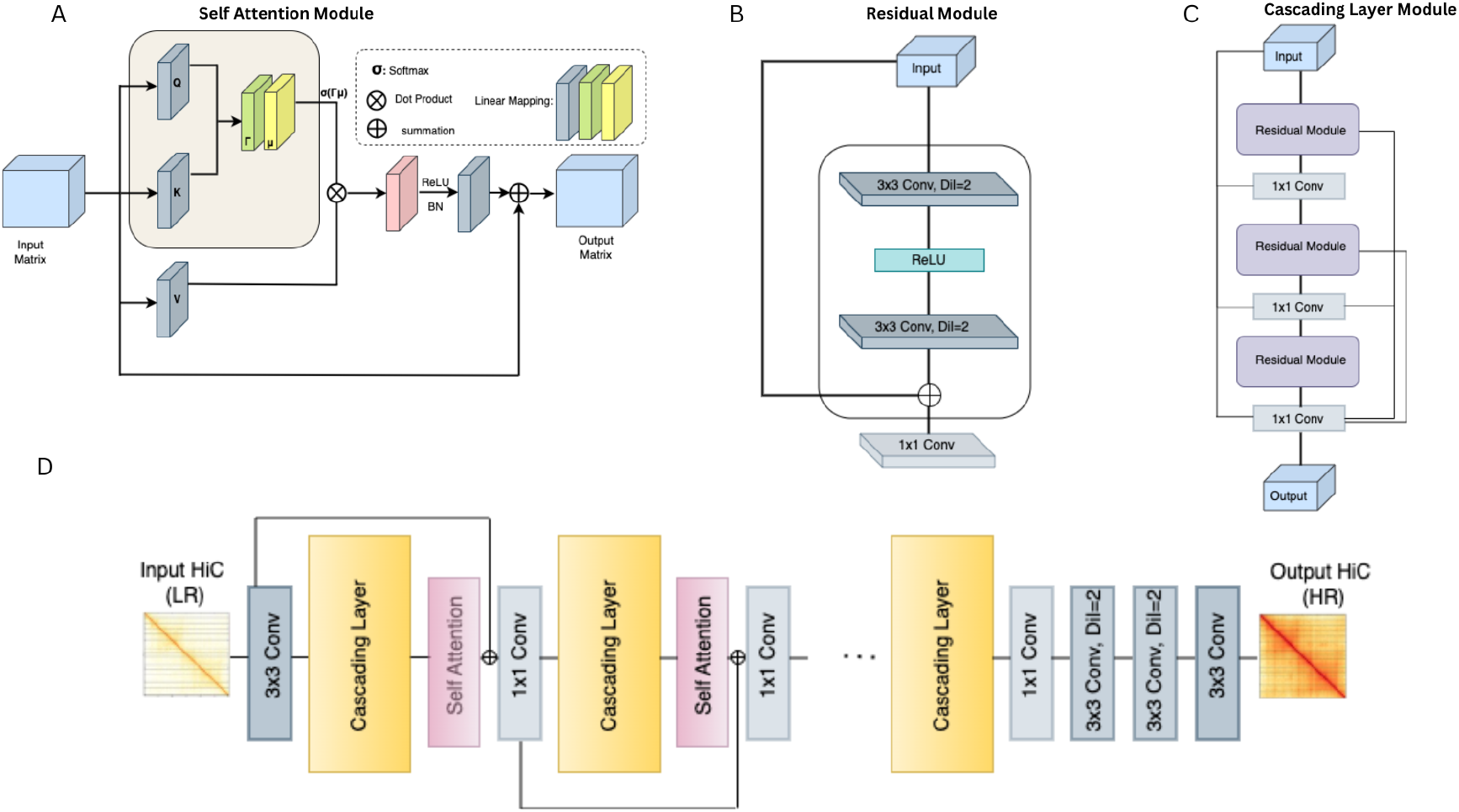
The DiCARN architecture comprises four major components. A shows the self-attention module that follows every instance of the cascading layer. B elucidates the residual module, which includes two dilated 3×3 convolutions, a ReLU activation function, a skip connection, and the 1×1 convolution in the concluding part. C outlines the cascading layer, which encapsulates the residual modules separated by 1×1 convolutions

#### 2.1.1 Dilation

Typically, the kernels in a convolution are contiguous. Dilation follows the ‘a trous algorithm, a technique used to increase the receptive field of the convolution operation by spacing out the kernel points without incrementing the number of parameters or the filter size ((X. Zhang *et al*., 2015)). “Trous” is a French term for “with holes”, essentially describing the implementation of dilated convolutions as the inclusion of gaps in the vanilla convolution operation.

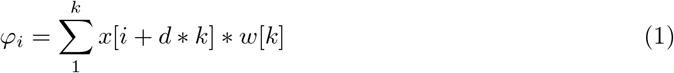

Equation 1 expresses this concept mathematically, where *φ*_*i*_ is the computed feature map, *d* is the dilation rate, *k* denotes kernel size, *w*[*k*] symbolizes the kernel weights, and *x*[*i*] signifies the input feature map.

We use dilated convolution within the residual module (Figure 1B) of our cascading layers and at the tail end of the entire network (Figure 1D) just before the final enhanced output is produced.

#### 2.1.2 Spatial Self-Attention

One of the long-standing challenges contending with CNN-based methods is their mode of treating all data point loci equally, thereby fostering redundancy in their computation of low-resolution features ((Q. Zhang *et al*., 2022), (Lu and X. Hu, 2022)). In our effort to redress this problem, we adopt the spatial self-attention method of Transformers ((Vaswani *et al*., 2017)).

The adoption of the spatial self-attention mechanism in our method (Figure 1A) is geared towards fostering a dynamic focus on different parts of the original LR feature map, ensuring a dynamic ability to model complex spatial dependencies and enhancing its ability to capture vital contextual information across spatial dimensions. More details are provided in Supplementary Section S1.

Utilizing the spatial self-attention mechanism in our method fosters a dynamic focus on different parts of the original LR feature map, ensuring a dynamic ability to model complex spatial dependencies and enhancing its ability to capture vital contextual information across spatial dimensions.

#### 2.1.3 Cascading ResNet

DiCARN employs a serialized cascade (Figure 1C) of multiple Residual Network (ResNet) modules (Ahn *et al*., 2018). Each residual module consists of two 3 × 3 convolutional layers, followed by a ReLU activation function, and incorporates a skip connection, enabling the network to retain information from earlier layers. This architectural design enhances the training process by mitigating issues such as vanishing gradients. Furthermore, feature representation is progressively refined in each cascading layer, as outputs from successive residual modules are combined to enhance the network’s overall performance.

### 2.2 Loss Function

The DiCARN model training employs the Mean Squared Error (MSE) loss function, leveraging its effectiveness in minimizing the difference between predicted and target values. This choice also aims to ensure computational simplicity and an exclusive focus on error minimization between predicted and observed Hi-C matrices. Equation 2 depicts the MSE, where *m* is the IF dimension, *P*_*a,b*_ and *Q*_*a,b*_ are the ground truth and predicted IFs between distal loci *a* and *b*, respectively.

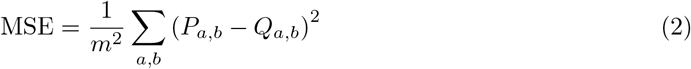

### 2.3 Data and Preprocessing

Our choice of Hi-C dataset is informed by the work of (Rao *et al*., 2014), which provides Hi-C data for several human cell types (K562, HMEC, NHEK, and GM12878), as well as the CH12-LX mouse cell line. All datasets are available in the NCBI GEO Accession Database under Accession ID: GSE63525. We trained our model using the GM12878 cell data, ensuring balance by excluding the X and Y chromosomes to avoid sex-related biases. Following the random chromosome selection method used in (Hicks and Oluwadare, 2022), validation was performed using chromosomes 2, 6, 10, and 12, while chromosomes 1 through 22, excluding chromosomes 4, 14, 16, and 20, were used for training. These excluded chromosomes were later utilized across cell lines for testing our model. For usability ease, all data used was split into small blocks of 40×40 dimensions.

### 2.4 Evaluation Metrics

To ensure a fair comparison, we adopt the Structural Similarity Index Measure (SSIM) and the Peak Signal-to-Noise Ratio (PSNR), the two favored computational metrics used in this research domain (Hong *et al*., 2020). We also used GenomeDISCO (Ursu *et al*., 2018) for our concordance measure and HiCRep (Yang *et al*., 2017) for the assessment of biological reproducibility. The SSIM between two images *x* and *y* is mathematically expressed as shown in Equation (3) where *µ*_*x*_ and *µ*_*y*_ are the mean intensities of images

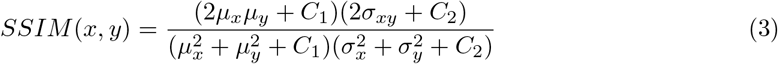

*x* and *y*, 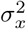 and 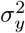 are the variances of images *x* and *y, σ*_*xy*_ is the covariance of images *x* and *y, C*_1_ and *C*_2_ are stability constants. Equation (4) gives the mathematical representation of the PSNR between two images. *L* is the maximum possible

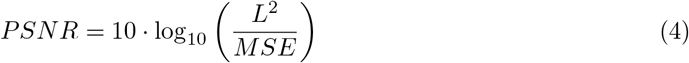

pixel value of the image (e.g., 255 for 8-bit images), while *MSE* is the Mean Squared Error between the given images.

Equation 5 shows the mathematical derivation of the concordance score as proffered by GenomeDISCO

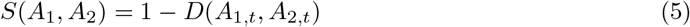

The concordance score given by this formula ranges between −1 and 1, where higher values signify greater similarity between the subject contact maps. Given two denoised contact maps *A*_1,*t*_ and *A*_2,*t*_, the difference between them is calculated using the distance *L*_1_ as shown in Equation (6).

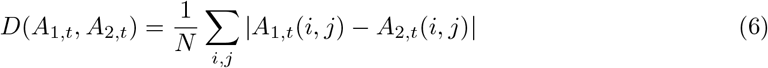

HiCRep, whose computation is presented in Equation (7) is the measure of the stratum-adjusted correlation coefficient (SCC) where *X*_*k*_ and *Y*_*k*_ are the contact frequencies contained in the stratum *k*, cov(*X*_*k*_, *Y*_*k*_) is the measure of covariance

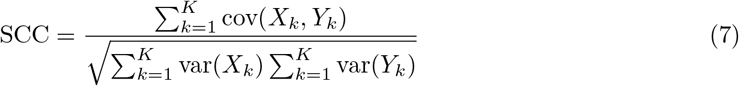

between *X*_*k*_ and *Y*_*k*_, var(*X*_*k*_) and var(*Y*_*k*_) are the variances of *X*_*k*_ and *Y*_*k*_ within every stratum, *K* is the sum total of strata.

## 3 Results

### 3.1 Hyperparameter search

Our proposed model is hinged on three key hyperparameters. (1) Number of Cascading Layers: HiCARN (Hicks and Oluwadare, 2022) performed a detailed hyperparameter search to determine the optimal number of cascading blocks in their work. They found that five cascading blocks provided the optimal result. Hence, we adopted the same number of blocks. (2) Self-Attention: To determine how to incorporate self-attention, we experimented with different configurations and the application of self-attention in different layers of our cascade architecture. Our optimal model was obtained by applying spatial self-attention to only the first two cascade blocks (Figure 1D), as shown in Supplementary Table S1. (3) Dilation Rate: To determine the dilation rate, we performed a hyperparameter search across dilation rates 2 to 5 and configurations. Our results show that dilation rate of 2 - with a configuration involving two dilated convolutions in the residual block as featured in Figure 1B and a dilated convolution stack at the end of the network, Figure 1D produced the optimal result (Supplementary Tables S2 and S3).

### 3.2 Training, Validation, and Testing

In the training phase, we conduct a validation after every training epoch so that the progressive performance of the model is accurately tracked and the optimally performing model weights are saved accordingly. This validation performance is then benchmarked against existing state-of-the-art methods (Figure 2). More results are presented in Supplementary Fig S1. The trainings were done using the low-resolution (LR) Hi-C dataset downsampled from the 10kb high-resolution variant made available in the GEO database accession number GSE63525. All models were trained on an NVIDIA GeForce RTX 4090 GPU with 24GB of VRAM, and the system had 128GB of RAM.

**Figure 2:**
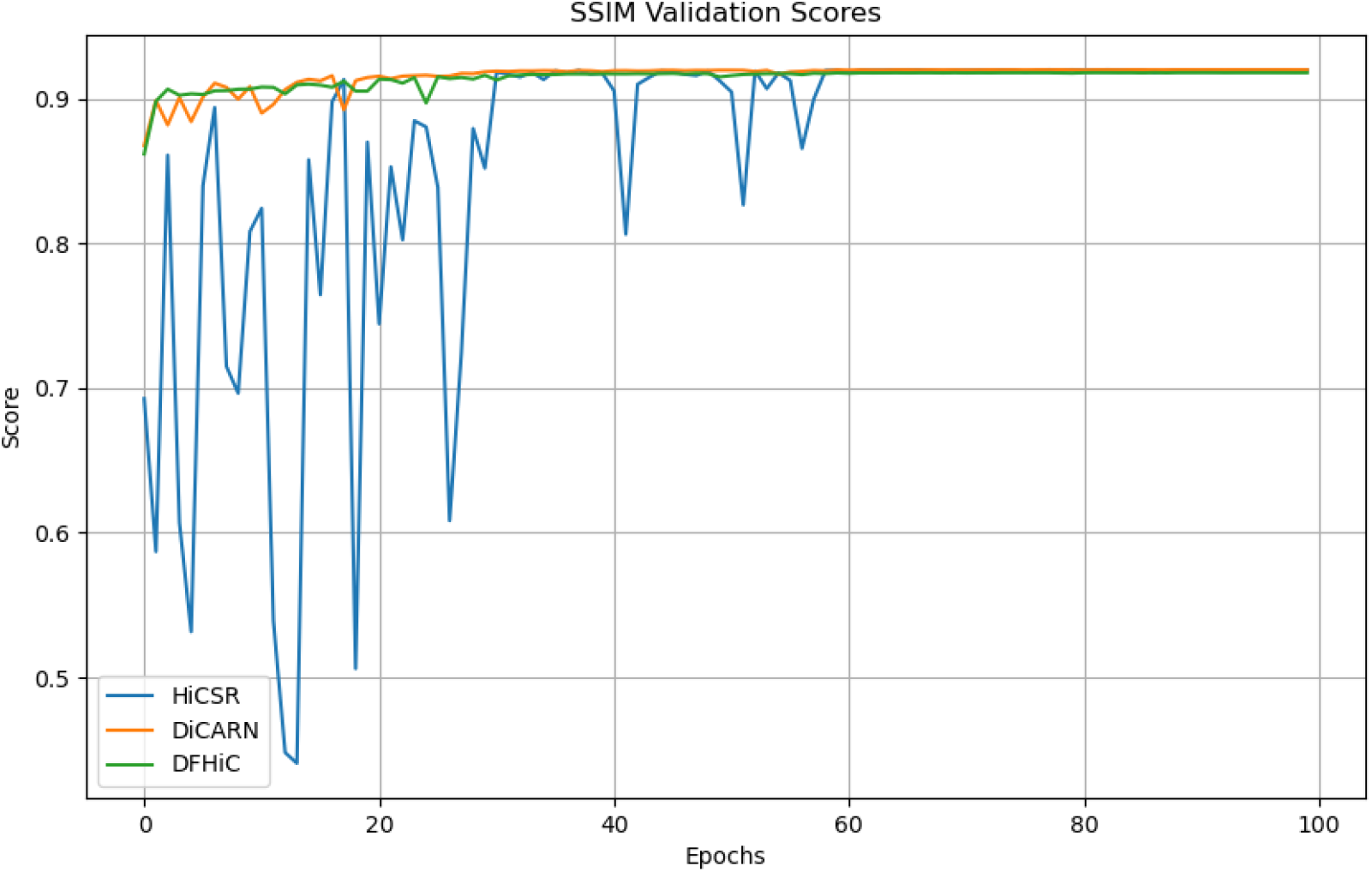
Validation result for DiCARN and state-of-the-art algorithms. Using the SSIM metric, the DiCARN validation results are contrasted with two state-of-the-art methods, HiCSR and DFHiC.

### 3.3 DiCARN Performance on Same Cell Line Data

HiCSR (Dimmick, 2020) and DFHiC (B. Wang *et al*., 2023) were selected for comparison with our model due to their demonstrated efficacy in GAN-based and CNN-based Hi-C data enhancement pipelines, respectively. After training our model on the 40kb low-resolution GM12878 Hi-C dataset, DiCARN consistently outperformed state-of-the-art models in both computational efficiency and biological benchmarks when tested on previously unseen GM12878 chromosomes. As shown in Table I, DiCARN’s same-cell prediction results exhibit superior performance relative to existing models. Additionally, we evaluated the model’s performance using a 1/64 downsample ratio, with the corresponding results provided in Supplementary Table S4. The training time and peak memory usage of the examined models are documented in Supplementary Figures S6 and S7, respectively.

**Table I:** DiCARN is benchmarked against existing methods based on same-cell GM12878 data with a downsampling ratio of 1/16 and it shows the highest performance on Average.

| Metrics | Method | Chr 4 | Chr 14 | Chr 16 | Chr 20 | Average |
| --- | --- | --- | --- | --- | --- | --- |
| SSIM | HiCSR | 0.93 | 0.9101 | 0.908 | 0.9086 | 0.9142 |
|  | DFHiC | 0.9292 | 0.9093 | 0.9055 | 0.9074 | 0.9129 |
|  | DiCARN | 0.9315 | 0.912 | 0.9111 | 0.9097 | **0.9161** |
| PSNR | HiCSR | 36.8908 | 35.446 | 33.9085 | 34.9226 | 35.292 |
|  | DFHiC | 36.7807 | 35.3461 | 33.609 | 34.7944 | 35.1326 |
|  | DiCARN | 36.9576 | 35.4686 | 33.9062 | 34.9067 | **35.3098** |
| Genome Disco | HiCSR | 0.9146 | 0.9219 | 0.904 | 0.9256 | 0.9165 |
|  | DFHiC | 0.9151 | 0.922 | 0.9045 | 0.9253 | 0.9167 |
|  | DiCARN | 0.9175 | 0.9243 | 0.9073 | 0.9273 | **0.9191** |
| HiCRep | HiCSR | 0.9179 | 0.8932 | 0.9408 | 0.8621 | 0.9035 |
|  | DFHiC | 0.9178 | 0.8928 | 0.9406 | 0.862 | 0.9033 |
|  | DiCARN | 0.9188 | 0.8928 | 0.9409 | 0.8622 | **0.9037** |

### 3.4 DiCARN Generalizability Test Across Unseen Cell Lines

Having trained the models on GM12878 cell type data only, we show in Table II that DiCARN generalizes better than existing state-of-the-art methods on the Lymphoblast cell line (K562), the Mammary Epithelial cell line (HMEC), and the human Epidermal Keratinocytes cell line (NHEK).

**Table II:** Performance benchmarking of state-of-the-art methods in contrast with DiCARN across unseen cell lines using average scores on test data. The results suggest that DiCARN retains its ability to restore the fidelity of *in silico* Hi-C data from unseen cell lines.

| Cell Line | Method | SSIM | PSNR | MSE |
| --- | --- | --- | --- | --- |
| K562 | HiCSR | 0.9485 | 34.7512 | **0.0003** |
|  | DFHiC | 0.9472 | 34.1669 | **0.0003** |
|  | DiCARN | **0.9498** | **35.2087** | **0.0003** |
| HMEC | HiCSR | 0.9748 | 35.2839 | **0.0002** |
|  | DFHiC | 0.9737 | 34.7582 | 0.0003 |
|  | DiCARN | **0.9757** | **35.2237** | **0.0002** |
| NHEK | HiCSR | 0.9721 | 35.093 | **0.0002** |
|  | DFHiC | 0.9705 | 34.0983 | 0.0003 |
|  | DiCARN | **0.9739** | **35.4481** | **0.0002** |

The experiment in this phase is based on the 1/16 ratio downsampled datasets for training and testing. The results maintain that DiCARN retains superior potential to generalize to unseen cell lines. We also present a visualization comparison of the corresponding structure similarity index measure for chromosomes 4, 14, and 20 for the different algorithms in Figure 3.

**Figure 3:**
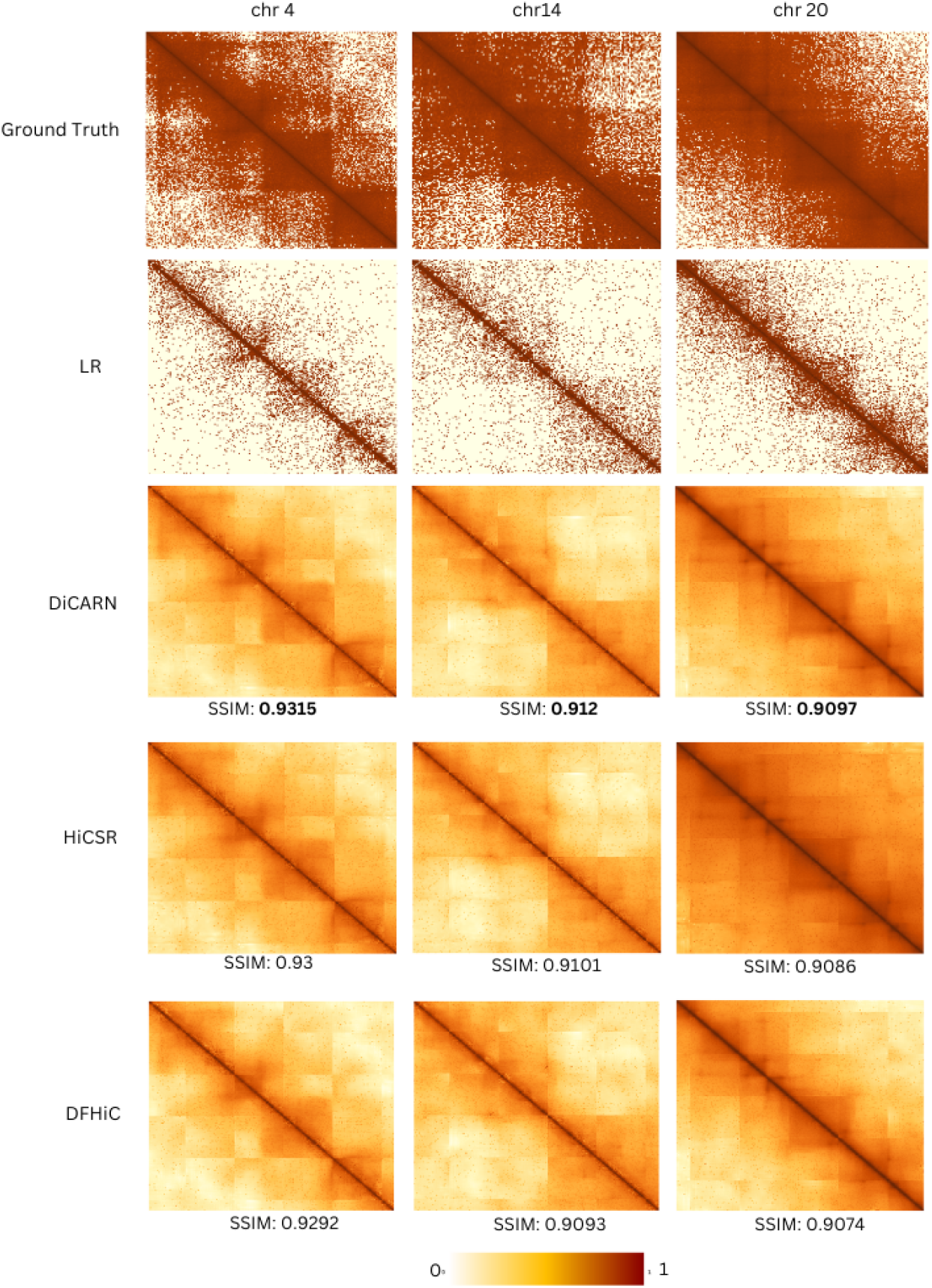
Heatmap visualization of the cross-cell enhancement results for DiCARN in comparison with state-of-the-art models (HiCSR and DFHiC) on chromosomes 4, 14, and 20 of the K562 cell line.

### 3.5 Enhancing Generalizability with Chromatin Accessibility Data from DNase-seq

#### 3.5.1 3.5.1. Chromatin Accessibility - DNase-seq Data

DNase-seq data denotes cell-type-specific chromatin accessibility and is a crucial marker in assessing 3D spatial organization due to its unique association with genomic regulatory elements (Yueqi Qiu *et al*., 2023). A recent application of chromatin accessibility data by Wang et al. (H. Wang *et al*., 2022) proposed a linear regression model to impute 3D distances between loci using DNase-seq data, thereby enhancing high-resolution 3D genome reconstruction accuracy.

Building upon this, we propose a novel approach for high-resolution Hi-C enhancement that leverages DNase-seq data to address the limitations of conventional Hi-C enhancement algorithms. We derive interaction frequencies (IF) from the DNase-seq and ultimately utilize this data to augment our training set and improve the generalizability of our model. The IF derived from DNase-seq is cell-type specific and is expected to enable accurate predictions across different biological contexts.

To calculate asynchronous interaction frequencies from DNase-seq, we employ a linear regression model, as shown in Equation (8) (H. Wang *et al*., 2022).

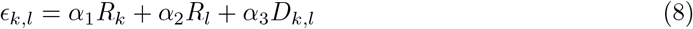

Our DNase-based IF imputation procedure, which enhances the resolution of 3D genomic maps, begins with the normalization of raw Hi-C interaction counts using Knight-Ruiz (KR) normalization to produce a normalized interaction frequency matrix. This matrix is then symmetrized to maintain consistency, and a Pairwise Distance (PD) matrix is generated to reflect spatial proximities among genomic loci. Due to the size of the interaction matrices, we fragment the data into manageable chunks, mapping each chunk to the corresponding DNase-seq signal using *bedtools* ((Quinlan and Hall, 2010)), thereby aligning chromatin accessibility data with the genomic coordinates. The DNase signal across each fragment is averaged to provide a summary measure of chromatin accessibility for each genomic region. Using these genomic distances and DNase signals as input, we predict distances using the pre-trained model defined in Equation (8) where *ϵ*_*k,l*_ represents the predicted interaction frequency between fragments *k* and *l, R*_*k*_ and *R*_*l*_ denote the DNase-seq signal levels for fragments *k* and *l, D*_*k,l*_ is the 1D genomic distance between these fragments, while *α*_1_, *α*_2_, and *α*_3_ are fitting parameters derived from imputed 3D distances ((H. Wang *et al*., 2022)). Subsequently, the imputed distances are converted into interaction frequencies (Lieberman-Aiden *et al*., 2009). The final reassembly process involves combining these IF matrix fragments into a complete matrix for the chromosome, followed by a KR normalization to ensure consistency with the original Hi-C data. Ultimately, this process produces a DNase-inferred IF matrix that supplements Hi-C data to refine resolution and improve interpretability across diverse cell types.

#### 3.5.2 Improving Generalizability Across Unseen Cell Lines with DNase-seq Data

To further enhance the generalizability of our model, we incorporate DNase-inferred IF data into the existing training dataset through a targeted data augmentation strategy intended to bolster the model’s predictive capacity across various cell lines. This strategy is explored in two configurations. In the target DNase scenario, DNase-imputed interaction frequency data from the target cell line’s test chromosomes is appended to the original Hi-C training dataset, after which the augmented dataset is used to retrain the DiCARN model. This model is called DiCARN-DNase-T(Supplementary Fig. S2). Specifically, these chromosomes correspond to those downsampled for testing. *We hypothesize that DNase-seq inferred IF data from the target cell type provides valuable insights into Hi-C interactions, facilitating enhanced model generalizability across varying biological contexts*.

The source DNase scenario, on the other hand, utilizes DNase-inferred IF data from the source cell type’s training chromosomes and adds it to the source dataset for training. This model is called DiCARN-DNase-S (Supplementary Fig. S3). *We hypothesize that including source DNase-based data not only augments the training set but also endows the model with a deeper understanding of chromatin dynamics beyond the source cell type*.

In this study, the source dataset is the GM12878 Hi-C dataset, and the target cell is the cell line on which we are testing the generalization.

Given that DNase data is expected to improve biological reproducibility, we evaluate our model’s performance using Hi-C analysis metrics the Stratum-adjusted Correlation Coefficient (SCC) through HiCRep, and the concordance score by GenomeDISCO. These metrics provide more biologically significant analysis measures compared to standard image evaluation metrics.

### 3.6 DiCARN-DNase: Enhancing DiCARN with DNase-seq for Cross-Cell Line Generalization

Table III highlights the HiCRep scores obtained for both configurations (DiCARN-DNase-T and DiCARN-DNase-S), where we demonstrate that at least one implementation of the DNase-augmented models outperforms the vanilla DiCARN model. Additionally, the results indicate that both augmentation strategies are viable, as each shows improvement over the vanilla model labelled DiCARN in at least one instance, thereby supporting our hypothesis. Our study also shows that there is no clear preference for one over the other as they could be viable options for both. The GenomeDISCO scores are presented in Supplementary Table S5 and Fig. S4.

**Table III:** HiCRep Average Score Comparison of DiCARN and its DNase-based variants on 1/16 downsampled K562 Cell Line dataset.

| Cell | Method | Chr 4 | Chr 14 | Chr 16 | Chr 20 | Avg |
| --- | --- | --- | --- | --- | --- | --- |
| K562 | DiCARN | 0.893 | 0.8466 | 0.9354 | 0.8478 | 0.8807 |
|  | DiCARN-DNase-S | 0.8914 | 0.8452 | 0.9334 | 0.8469 | 0.8792 |
|  | DiCARN-DNase-T | 0.8931 | 0.8466 | 0.9417 | 0.8465 | **0.884** |
| HMEC | DiCARN | 0.7302 | 0.7794 | 0.8193 | 0.6986 | 0.7570 |
|  | DiCARN-DNase-S | 0.7181 | 0.7716 | 0.8140 | 0.6923 | 0.7490 |
|  | DiCARN-DNase-T | 0.7223 | 0.7752 | 0.8187 | 0.7132 | **0.7574** |
| NHEK | DiCARN | 0.8161 | 0.8158 | 0.8493 | 0.7217 | 0.8007 |
|  | DiCARN-DNase-S | 0.8172 | 0.8153 | 0.8482 | 0.7293 | **0.8012** |
|  | DiCARN-DNase-T | 0.8195 | 0.8196 | 0.8516 | 0.7312 | **0.8055** |

### 3.7 Enhancing Generalizability of State-of-the-art Models with DNase-seq Data

Following the enhancement capability boost recorded by DiCARN when influenced by the IF imputed from the DNase-seq data, we proceeded to appropriate this data augmentation innovation to some existing models in the Hi-C resolution enhancement research domain, including HiCSR (Dimmick, 2020), HiCARN (Hicks and Oluwadare, 2022), HiCNN(Liu and Z. Wang, 2019), and DFHiC (B. Wang *et al*., 2023) to test the generalizability of the DNase idea to other models. From the HiCRep results presented in Table IV, it is observed that the data augmentation approach worked in ten of twelve scenarios. More results are provided in Supplementary Table S6 and Fig. S5. It is also observed that the DFHiC was the base method in the two instances where the approach was challenged. However, the majority of results obtained from the test for applicability to other methods established the proposition that the fusion of DNase with *in silico* LR GM12878 Hi-C data improves the cell-to-cell Hi-C reproducibility capabilities of deep learning-based methods. This exception leads us to believe that the data augmentation might be sensitive to the algorithm of the method.

**Table IV:** Using HiCRep for reproducibility assessment, we show the performance scores of the DNase-based data augmentation approach for four existing DL methods across three cell lines. Each vanilla method is contrasted with its corresponding Target DNase (e.g. HiCSR-DNase-T) and Source DNase (e.g. HiCSR-DNase-S) variants. We observe that the majority of the test cases support the DNase proposition.

| Cell | Method | Chr 4 | Chr 14 | Chr 16 | Chr 20 | Avg |
| --- | --- | --- | --- | --- | --- | --- |
| K562 | HiCSR | 0.8913 | 0.8444 | 0.9304 | 0.8414 | 0.8769 |
|  | HiCSR-DNase-S | 0.8929 | 0.8469 | 0.9295 | 0.8444 | **0.8784** |
|  | HiCSR-DNase-T | 0.8923 | 0.8445 | 0.9304 | 0.8441 | 0.8778 |
|  | DFHiC | 0.8905 | 0.8423 | 0.9338 | 0.8424 | **0.8773** |
|  | DFHiC-DNase-S | 0.8903 | 0.8424 | 0.9256 | 0.8333 | 0.8729 |
|  | DFHiC-DNase-T | 0.8869 | 0.8398 | 0.9307 | 0.8367 | 0.8735 |
|  | HiCARN | 0.8928 | 0.8444 | 0.9375 | 0.8496 | 0.8811 |
|  | HiCARN-DNase-S | 0.8968 | 0.8482 | 0.9289 | 0.8571 | **0.8828** |
|  | HiCARN-DNase-T | 0.8913 | 0.8424 | 0.9324 | 0.849 | 0.8788 |
|  | HiCNN | 0.8956 | 0.8477 | 0.9257 | 0.8378 | 0.8767 |
|  | HiCNN-DNase-S | 0.8946 | 0.8478 | 0.941 | 0.8538 | **0.8843** |
|  | HiCNN-DNase-T | 0.8903 | 0.8434 | 0.9359 | 0.847 | 0.8792 |
| HMEC | HiCSR | 0.7296 | 0.7793 | 0.8201 | 0.6998 | 0.7572 |
|  | HiCSR-DNase-S | 0.7223 | 0.7734 | 0.8155 | 0.6935 | 0.7512 |
|  | HiCSR-DNase-T | 0.74 | 0.7734 | 0.8554 | 0.694 | **0.7657** |
|  | DFHiC | 0.7328 | 0.7775 | 0.8212 | 0.6997 | 0.7578 |
|  | DFHiC-DNase-S | 0.7244 | 0.7791 | 0.8257 | 0.705 | **0.7586** |
|  | DFHiC-DNase-T | 0.738 | 0.7844 | 0.8244 | 0.7019 | **0.7622** |
|  | HiCARN | 0.7269 | 0.776 | 0.8191 | 0.6947 | 0.7542 |
|  | HiCARN-DNase-S | 0.7192 | 0.7693 | 0.8143 | 0.6926 | 0.7498 |
|  | HiCARN-DNase-T | 0.7312 | 0.7796 | 0.8218 | 0.7012 | **0.7585** |
|  | HiCNN | 0.7182 | 0.7711 | 0.8144 | 0.6952 | 0.7497 |
|  | HiCNN-DNase-S | 0.7182 | 0.7719 | 0.815 | 0.6966 | **0.7504** |
|  | HiCNN-DNase-T | 0.7257 | 0.7762 | 0.8163 | 0.6957 | **0.7535** |
| NHEK | HiCSR | 0.8165 | 0.8152 | 0.8498 | 0.7242 | 0.8014 |
|  | HiCSR-DNase-S | 0.8191 | 0.8169 | 0.8509 | 0.7282 | 0.8038 |
|  | HiCSR-DNase-T | 0.8194 | 0.817 | 0.8509 | 0.7283 | **0.8039** |
|  | DFHiC | 0.8168 | 0.8167 | 0.8504 | 0.7239 | **0.8029** |
|  | DFHiC-DNase-S | 0.8121 | 0.8126 | 0.8469 | 0.7189 | 0.7976 |
|  | DFHiC-DNase-T | 0.8131 | 0.8152 | 0.8487 | 0.7177 | 0.7987 |
|  | HiCARN | 0.8157 | 0.8136 | 0.8462 | 0.7229 | 0.7996 |
|  | HiCARN-DNase-S | 0.8123 | 0.8083 | 0.8405 | 0.7246 | 0.7964 |
|  | HiCARN-DNase-T | 0.8186 | 0.8163 | 0.8494 | 0.7266 | **0.8027** |
|  | HiCNN | 0.82 | 0.8159 | 0.8 | 0.73 | 0.7915 |
|  | HiCNN-DNase-S | 0.8223 | 0.8183 | 0.841 | 0.73 | **0.8029** |
|  | HiCNN-DNase-T | 0.8089 | 0.806 | 0.8392 | 0.717 | 0.7928 |

### 3.8 3.8. DiCARN-DNase: Leading Performance in Hi-C Data Enhancement Across Cell Lines

The integration of DNase-seq data significantly enhances the performance of both our model and existing algorithms. To assess the overall performance of the algorithms (including both vanilla and DNase-augmented variants) across different cell lines, we constructed a ranking table based on HiCRep scores, which measure consistency across three cell lines using the average results from four test chromosomes. The scores for DiCARN are presented in Table III, while the results for the other four algorithms are provided in Table IV. As shown in the ranking table (Table V), the DiCARN models demonstrated superior performance, achieving an average rank of 2.3. Specifically, DiCARN ranked first for NHEK, second for K562, and fourth for HMEC, indicating a high degree of generalizability across cell lines, based on the HiCRep scores.

**Table V:** Comparative Performance Rankings of DiCARN and Other Models Across Cell Lines. The number represent rank in terms of performance where a lower number indicates better performance (e.g., a ranking of 1 is better than 5)

| Cell | DiCARN | HiCSR | HiCARN | DFHiC | HiCNN |
| --- | --- | --- | --- | --- | --- |
| K562 | 2 | 4 | 3 | 5 | 1 |
| HMEC | 4 | 1 | 3 | 2 | 5 |
| NHEK | 1 | 4 | 5 | 2 | 2 |
| **Avg** | **2.3\*** | **3** | 3.66 | 3 | 2.66** |

### 3.9 3.9. Benchmark on 3D Genome Reconstruction and TAD Detection

The ability of the data from these models to recover Topologically Associating Domains (TADs) plays a critical role in exploring functional genomics and regulating gene expression by controlling enhancer-promoter interactions (Dixon *et al*., 2012) and also plays an important role towards usefulness. In this study, we employed TopDom (Shin *et al*., 2016) to detect TADs from region 60Kb to 2.45Mb region of K562 cell line chromosome 14 using the imputed Hi-C data and the ground truth data. We assessed their concordance through the Measure of Concordance (MoC) metric (Higgins *et al*., 2022). A higher MoC score is better. The results indicate that the DNase-based variants for most algorithms closely match the ground truth, underscoring the impact of DNase-seq data on enhancing Hi-C data (Figure 4).

**Figure 4:**
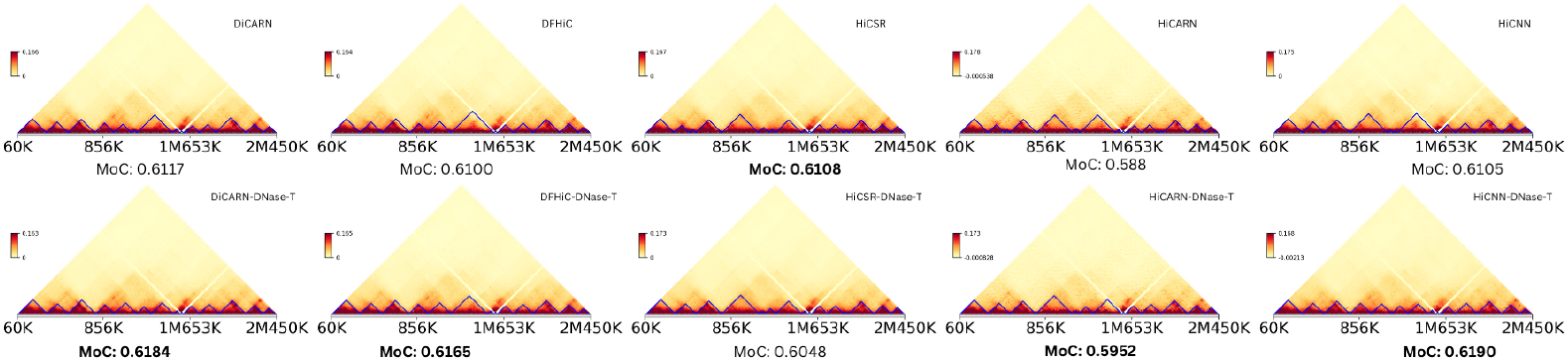
The TAD recovery assessment of the Baseline algorithm (Vanilla) compared versus the DNase-based variants. We used the 60Kb to 2.45Mb region of the chr14 in the K562 cell line for this procedure. The procedure was also executed across other methods to show their TAD recovery abilities. The heatmaps are tagged with their corresponding MoCs to show the improvement of the DNase-based models on the vanilla variants.

Furthermore, we evaluated the structural similarity of the 3D genome reconstructed from both the imputed and ground truth data. Using 3DMax (Oluwadare *et al*., 2018), we reconstructed structures for region 300Mb to 350Mb of K562 cell line chromosome 20 and compared them via the Spearman Correlation Coefficient (Figure 5). The results demonstrate that the DNase-augmented DiCARN model showed greater concordance with the ground truth than the vanilla DiCARN model. Overall, these findings affirm the potency of DNase-seq augmentation in the prediction accuracy of 3D genomic structures.

**Figure 5:**
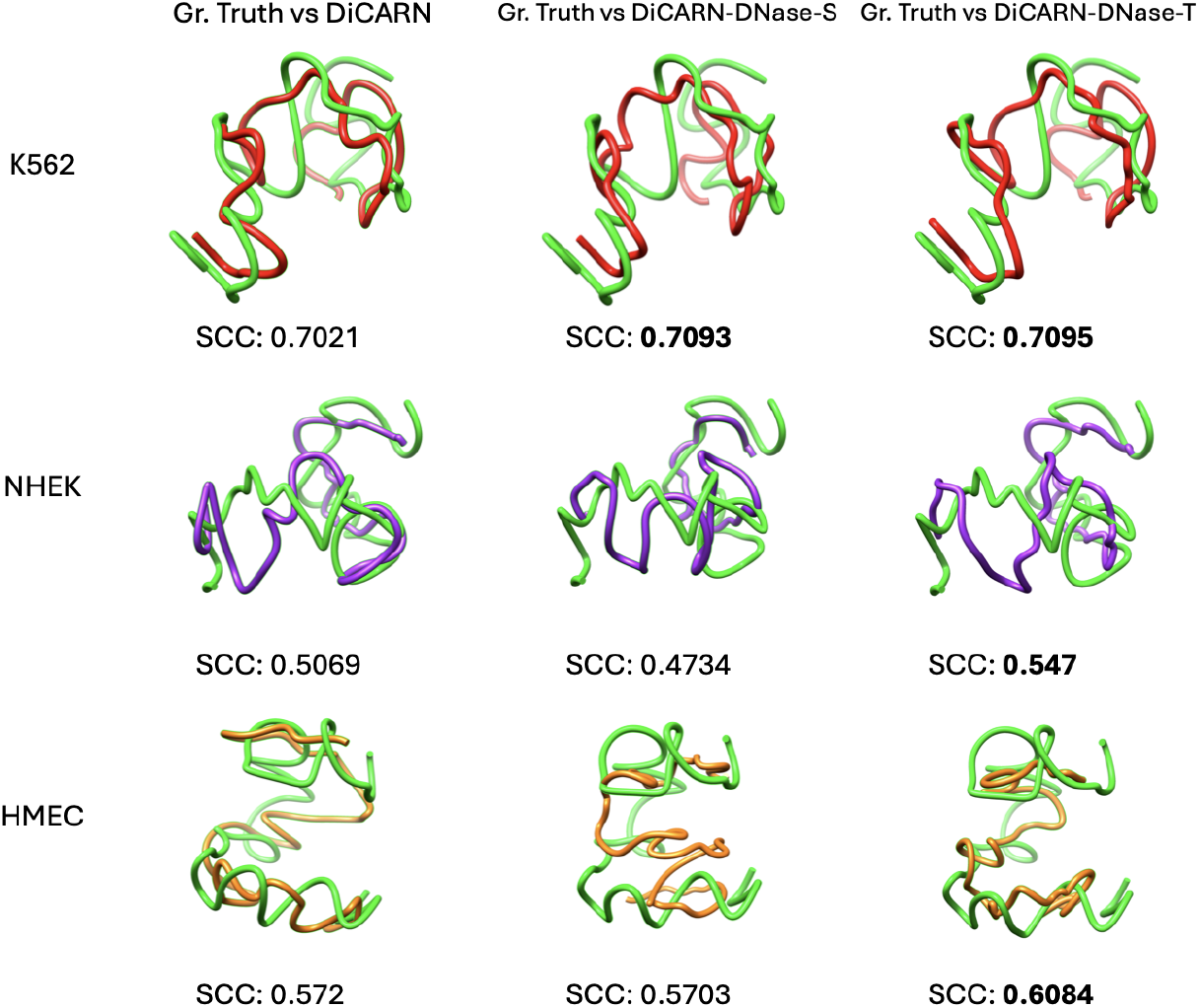
Evaluation of Groundtruth consistency with output from DiCARN variants for chromosome 20 of the K562 cell line using the 300MB to 350MB region. Reconstructed 3D structures for DNase-based DiCARN records more consistency based on the SCC scores than its vanilla variant.

## 4 Conclusion

In this study, we introduce DiCARN, an attention-based Dilated Cascading ResNet model for the recovery of high-resolution Hi-C data necessary for biological and computational exploits of genomic structures. Eminently, our study pivots on the introduction of a novel approach involving distal inferences from the chromatin accessibility DNase data of human cell lines for the augmentation of LR interaction frequency data. The practicality of this innovation was tested and established using biological reproducibility and structural similarity metrics. It is important to note that the inclusion of DNase-seq data has been universally beneficial across all models, including existing state-of-the-art models This study emphasizes how the use of DNase-seq data has elevated the performance of both our model and others.

## 5 Author contributions statement

S.O. designed the pipeline, wrote the code, and wrote the initial draft manuscript. S.O. and O.O. analyzed the results. O.O. conceived and supervised the project. All authors wrote and reviewed the manuscript.

## 6 Acknowledgements

I would like to thank Rohit Menon, HMA Choudhury, and Abhishek Pandeya for their resource contributions to this work.

## 7 Code and Data Availability

DiCARN-DNase is a containerized software made available via: https://github.com/OluwadareLab/DiCARN_DNase. The DNase-seq data was collected from Roadmap and Consortia Database (https://egg2.wustl.edu/roadmap/web_portal/processed_data.html), while the Hi-C datasets for the GM12878, K562, HMEC, and NHEK cell lines (Rao *et al*., 2014) used in this study are available in the GEO Accession Database via GEO code GSE63525.

## 8 Supplemental Data

Supplementary figures and tables are included in the Supplementary Materials document.

