## Supplemental Document for "DiCARN-DNase: Enhancing Cell-to-Cell Hi-C Resolution Using Dilated Cascading ResNet with Self-Attention and DNase-seq Chromatin Accessibility Data"

SUPPLEMENTARY DATA

### 1 The DiCARN Architecture

#### 1.1 Spatial Self Attention

Suppose that Hi-C feature map $B_i \subset \mathbb{R}^{HxWxD}$ (where i = q, k, v) is obtained post-convolutions from the spatial attention module input $B \subset \mathbb{R}^{HxWxD}$, denoting Query, Key, and Value respectively, we represent spatial attention operation as follows:

$$\mathbf{B}_i = \mathbf{G}_i(\mathbf{B}), \quad i = q, k, v \tag{1}$$

where $C(.)$ represents the convolution, which results in a C-dimensional feature map matrix $B_i$, a feature map featuring C channels. $B_i$ is then reshaped into a 2D form, expressed thus:

$$\mathbf{B}_{\text{reshape},i} = \mathbf{H}(\mathbf{B}_i), \quad i = q, k, v \tag{2}$$

After symmetrizing the query, key, and value outputs using the reshape function $H$, we follow up with a transpose and multiplication computation expressed as follows:

$$(\mathbf{B}_{\text{reshape},q})^T = \mathbf{U}(\mathbf{B}_{\text{reshape},q}) \tag{3}$$

Here, the $U$ represents the transpose mapping. This operation is then followed by the computation of the weight coefficient matrix M expressed thus:

$$\mathbf{M} = (\mathbf{B}_{\text{reshape},1})^T \times \mathbf{B}_{\text{reshape},2} \tag{4}$$

Softmax is now applied to the obtained weight coefficient matrix $M$ so that $M_{softmax}$ is then obtained:

$$b_{ij} = \frac{\exp(y_{ij})}{\sum_{k=1,l=1}^{N} \exp(y_{ij})}, \quad \mathbf{M}_{\text{softmax}} = [b_{ij}] \tag{5}$$

Finally, the attention feature map is computed as follows:

$$\mathbf{F} = \mathbf{B} + \beta \cdot \mathbf{U}(\mathbf{U}(\mathbf{M}_{\text{softmax}}) \times \mathbf{B}_{\text{reshape},3}) \tag{6}$$

Table 1: Average Scores of Attention on DiCARN Cascades

| **Attention on DiCARN Cascades** | | | |
|---|---|---|---|
| **Average Scores** | **SSIM** | **PSNR** | **GenomeDISCO** |
| Att. 1 Cascade | 0.9468 | 35.2252 | 0.8558 |
| Att. 2 Cascades | 0.9471 | **35.2986** | **0.8579** |
| Att. 3 Cascades | 0.9459 | 34.4689 | 0.8266 |
| Att. 4 Cascades | **0.9474** | 35.1339 | 0.8563 |
| Att. 5 Cascades | **0.9474** | 35.1815 | 0.8561 |

##### 1.2 Hyperparameter Search

Table 1 reports our search for the optimal application of spatial self-attention in our architectural pipeline. The results reported informed our decision to fit the DiCARN model with spatial self-attention on the first 2 cascades.

The decision on the number of dilations that work optimally for our model was determined by a series of tests. The results shown in Table 2 informed our decision to use the dilation rate of 2 in every instance of dilated convolution within the architecture of our model.

As shown in Table 3, our decision to append two 3x3 convolutions with a dilation rate of 2 is informed by the results recorded in this phase of our ablation study. Based on the performance observations at varying instances of dilation application on the tail of our model architecture, we adopt a dual-layer convolution with a dilation rate of 2 in our final structure.

Table 2: Performance metrics at various dilation rates within the residual blocks of the DiCARN model network.

| CHR | SSIM | PSNR | GenomeDISCO | Loss |
|---|---|---|---|---|
| | | Dilation Rate = 2 | | |
| 3 | 0.9532 | 34.1225 | 0.8478 | 0.0004 |
| 11 | 0.9472 | 34.3523 | 0.8564 | 0.0003 |
| 19 | 0.9508 | 35.8801 | 0.8567 | 0.0003 |
| 21 | 0.9327 | 34.008 | 0.8677 | 0.0004 |
| Average | 0.946 | 34.5907 | **0.8572** | **0.0003** |
| | | Dilation Rate = 3 | | |
| 3 | 0.954 | 34.2952 | 0.8484 | 0.0004 |
| 11 | 0.9481 | 34.7978 | 0.8564 | 0.0003 |
| 19 | 0.951 | 36.1661 | 0.8564 | 0.0003 |
| 21 | 0.9333 | 34.4736 | 0.8669 | 0.0004 |
| Average | **0.9466** | 34.4736 | 0.857 | **0.0003** |
| | | Dilation Rate = 4 | | |
| 3 | 0.9538 | 34.464 | 0.8474 | 0.0004 |
| 11 | 0.9477 | 34.8228 | 0.8559 | 0.0003 |
| 19 | 0.9508 | 36.1773 | 0.8554 | 0.0003 |
| 21 | 0.9327 | 34.3966 | 0.8676 | 0.0004 |
| Average | 0.9463 | **34.9652** | 0.8567 | **0.0003** |
| | | Dilation Rate = 5 | | |
| 3 | 0.9372 | 34.4214 | 0.8193 | 0.0004 |
| 11 | 0.9304 | 34.1406 | 0.8283 | 0.0004 |
| 19 | 0.9331 | 33.4715 | 0.8283 | 0.0005 |
| 21 | 0.9116 | 33.4666 | 0.8393 | 0.0006 |
| Average | 0.9281 | 33.875 | 0.8288 | 0.0005 |

#### 2 Results

##### 2.1 DiCARN Same Cell Results on GM12878 1/64 ratio Downsampled Data

To further establish our hypothesis about the performance of DiCARN, we repeat the high-resolution data prediction experiment, this time training on 1/64 downsampled Hi-C data of all but four GM12878, which were used for testing. This result is presented in Table 4.

##### 2.2 Concordance Comparison of TAD Recoverability Between Vanilla Models and Their DNase-Inspired Variants

Here we provide the concordance score comparison between five selected models and their corresponding DNase-based versions.

Table 3: Performance Metrics for DiCARN Models with varying dilated convolutional layers at the tail end of the architecture. *DiCARN + (2 X DL - Tail)* means the tail of the network comprises a stack of 2 dilated convolutions, etc.

| **CHR** | **SSIM** | **PSNR** | **GenomeDISCO** |
|---|---|---|---|
| DiCARN | | | |
| 3 | 0.9532 | 34.1225 | 0.8478 |
| 11 | 0.9472 | 34.3523 | 0.8564 |
| 19 | 0.9508 | 35.8801 | 0.8567 |
| 21 | 0.9327 | 34.008 | 0.8677 |
| Average | 0.946 | 34.5907 | 0.8572 |
| DiCARN + (2 x DL-Tail) | | | |
| 3 | 0.9535 | 34.4711 | 0.8473 |
| 11 | 0.9479 | 34.8474 | 0.8555 |
| 19 | 0.9514 | 36.152 | 0.8558 |
| 21 | 0.9337 | 34.4538 | 0.866 |
| Average | 0.9466 | 34.9811 | 0.856 |
| DiCARN + (3 x DL-Tail) | | | |
| 3 | 0.9536 | 34.5774 | 0.8396 |
| 11 | 0.9478 | 34.7324 | 0.8515 |
| 19 | 0.9513 | 35.7523 | 0.8516 |
| 21 | 0.9333 | 34.3046 | 0.8627 |
| Average | 0.9465 | 34.841675 | 0.8514 |
| DiCARN + (5 x DL-Tail) | | | |
| 3 | 0.9527 | 34.181 | 0.8448 |
| 11 | 0.9472 | 34.5606 | 0.855 |
| 19 | 0.9511 | 35.7801 | 0.8549 |
| 21 | 0.9329 | 34.1898 | 0.8667 |
| Average | 0.946 | 34.6779 | 0.8554 |

Table 4: Comparison of same cell SSIM, PSNR, GenomeDISCO and HiCRep scores across four different chromosomes basing the experimentation on the 1/64 ratio downsampled GM12878 datasets.

| **Method** | **Chromosome** | **SSIM** | **PSNR** | **GenomeDISCO** | **HiCRep** |
|---|---|---|---|---|---|
| HiCSR | Chr 4 | 0.9143 | 35.6552 | 0.8985 | 0.6758 |
| | Chr 14 | 0.8908 | 34.1203 | 0.9056 | 0.7477 |
| | Chr 16 | 0.8709 | 31.6647 | 0.8797 | 0.8497 |
| | Chr 20 | 0.8898 | 33.4203 | 0.9081 | 0.6697 |
| | Averages | 0.8915 | **33.7151** | 0.898 | 0.7357 |
| DFHiC | Chr 4 | 0.9106 | 35.2824 | 0.8932 | 0.6746 |
| | Chr 14 | 0.8873 | 33.8337 | 0.901 | 0.7458 |
| | Chr 16 | 0.864 | 31.0145 | 0.8753 | 0.8491 |
| | Chr 20 | 0.8854 | 33.0785 | 0.9038 | 0.669 |
| | Averages | 0.8868 | 33.3023 | 0.8933 | 0.7346 |
| DiCARN | Chr 4 | 0.9148 | 35.5979 | 0.8983 | 0.6783 |
| | Chr 14 | 0.8909 | 34.1 | 0.9055 | 0.7483 |
| | Chr 16 | 0.871 | 31.51 | 0.8795 | 0.8504 |
| | Chr 20 | 0.8897 | 33.362 | 0.91 | 0.6695 |
| | Averages | **0.8916** | 33.6425 | **0.8983** | **0.7366** |

Table 5: GenomeDISCO Average Score Comparison of DiCARN and its DNase-ased variants on 1/16 downsampled K562 Cell Line dataset

| **GenomeDISCO** | **Avg** |
|---|---|
| DiCARN | 0.8345 |
| DiCARN-DNase-S | 0.8364 |
| DiCARN-DNase-T | 0.8347 |

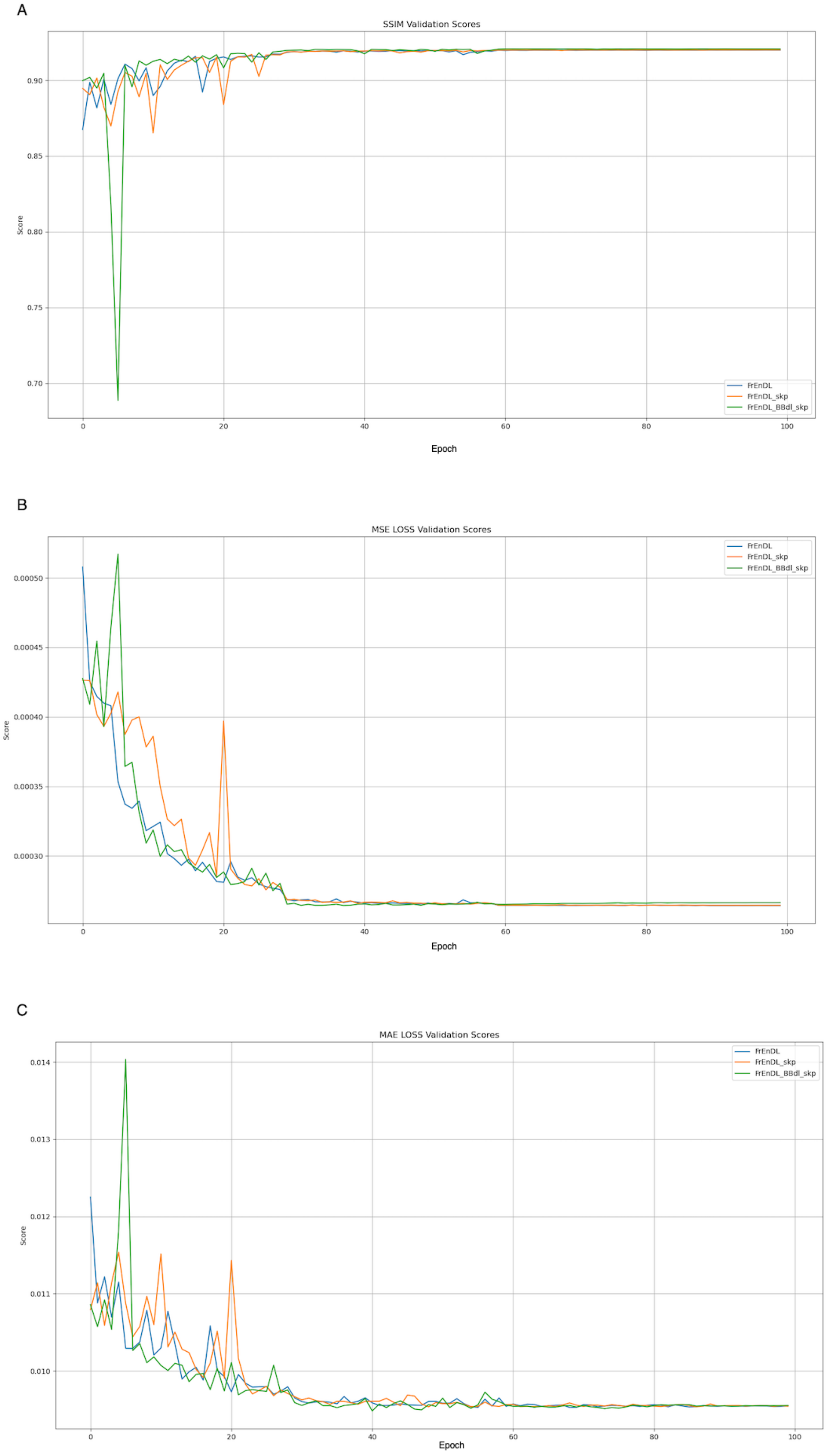


Figure 1: Validation plots for the structural decision three skip connection implementation variations. Figure A gives the SSIM score plot, plot B shows the comparison of the MSE Loss scores, and plot C depicts the MAE loss comparison between the three variants. FrEnDL is the DiCARN architectural variant that features no skip connections; FrEnDL-skp is the version featuring a moderate amount of skip connections within each cascade layer and between cascade layers; FrEnDL-BBdl-skp is the version that highlights an additional skip connection linking the positional embeddings obtained from the first basic block layer to the final 1x1 convolutional layer at the base of the network. This experiment shows that the FrEnDL works best and was adopted as the final DiCARN version.

Table 6: Comparison of GenomeDISCO for different models and their DNase-based counterparts across various chromosomes of the 1/16 downsampled K562 cell line.

| Method | Chromosome | GenomeDISCO |
|---|---|---|
| **DiCARN** | Chr 4 | 0.845 |
| | Chr 14 | 0.8122 |
| | Chr 16 | 0.8252 |
| | Chr 20 | 0.8557 |
| **DiCARN DNase Src** | Chr 4 | 0.8458 |
| | Chr 14 | 0.8141 |
| | Chr 16 | 0.8286 |
| | Chr 20 | 0.857 |
| **DiCARN DNase Target** | Chr 4 | 0.8444 |
| | Chr 14 | 0.8114 |
| | Chr 16 | 0.8267 |
| | Chr 20 | 0.8564 |
| **HiCSR** | Chr 4 | 0.839 |
| | Chr 14 | 0.805 |
| | Chr 16 | 0.8206 |
| | Chr 20 | 0.8501 |
| **HiCSR-DNase-Src** | Chr 4 | 0.8417 |
| | Chr 14 | 0.8087 |
| | Chr 16 | 0.8221 |
| | Chr 20 | 0.8522 |
| **HiCSR-DNase-Target** | Chr 4 | 0.8417 |
| | Chr 14 | 0.8092 |
| | Chr 16 | 0.8235 |
| | Chr 20 | 0.8525 |
| **HiCNN** | Chr 4 | 0.844 |
| | Chr 14 | 0.8124 |
| | Chr 16 | 0.8253 |
| | Chr 20 | 0.8554 |
| **HiCNN-DNase-Src** | Chr 4 | 0.8436 |
| | Chr 14 | 0.8111 |
| | Chr 16 | 0.859 |
| | Chr 20 | 0.8552 |
| **HiCNN-DNase-Target** | Chr 4 | 0.8427 |
| | Chr 14 | 0.8093 |
| | Chr 16 | 0.8248 |
| | Chr 20 | 0.8545 |
| **DFHiC** | Chr 4 | 0.844 |
| | Chr 14 | 0.8124 |
| | Chr 16 | 0.8253 |
| | Chr 20 | 0.8554 |
| **DFHiC-Dnase-Src** | Chr 4 | 0.8436 |
| | Chr 14 | 0.8111 |
| | Chr 16 | 0.859 |
| | Chr 20 | 0.8552 |
| **DFHiC-DNase-Target** | Chr 4 | 0.8427 |
| | Chr 14 | 0.8093 |
| | Chr 16 | 0.8248 |
| | Chr 20 | 0.8545 |

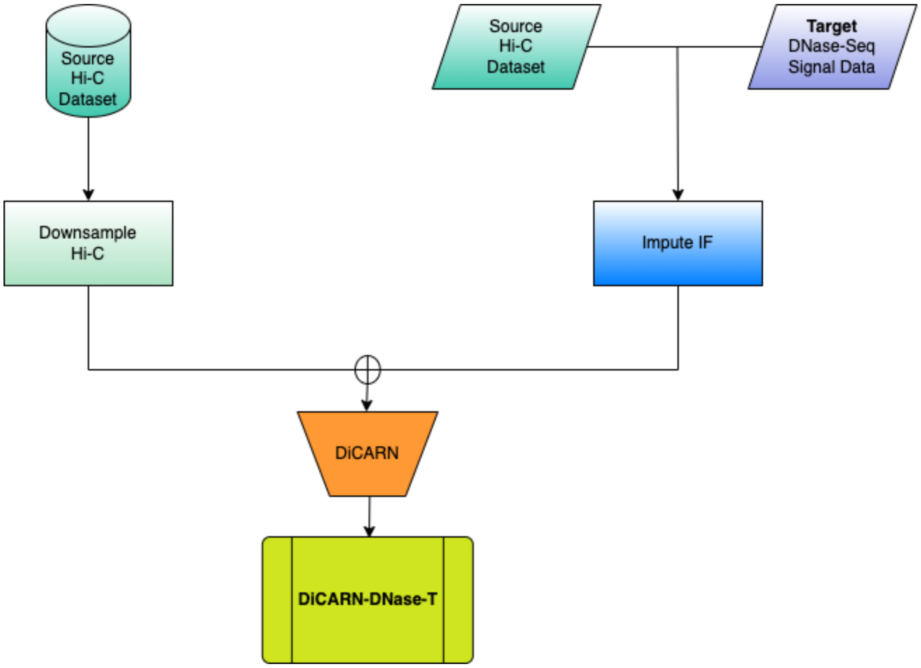


Figure 2: The pictorial illustration of the DiCARN-DNase-T

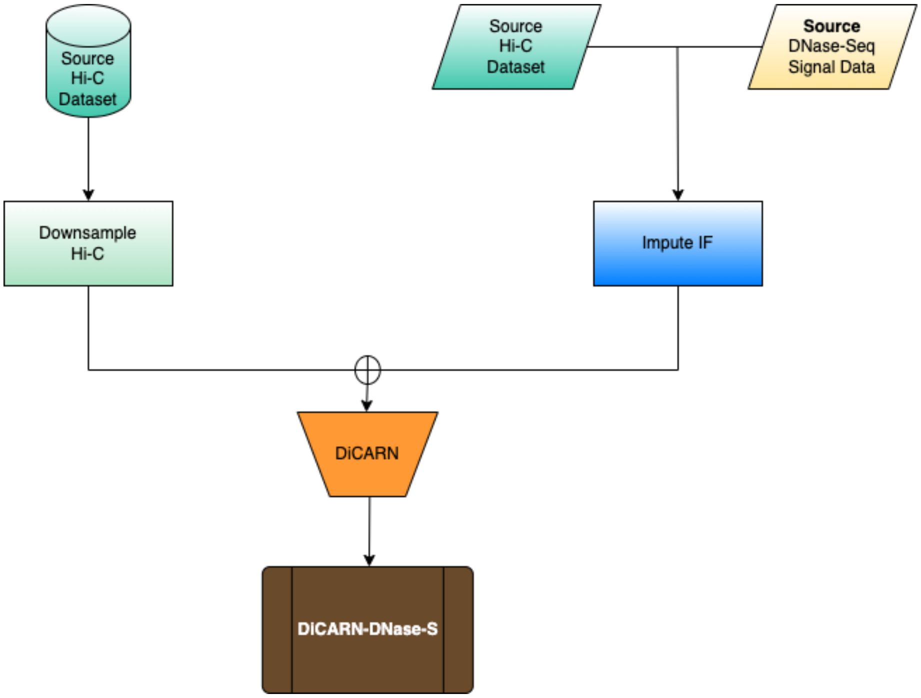


Figure 3: The pictorial illustration of the DiCARN-DNase-S

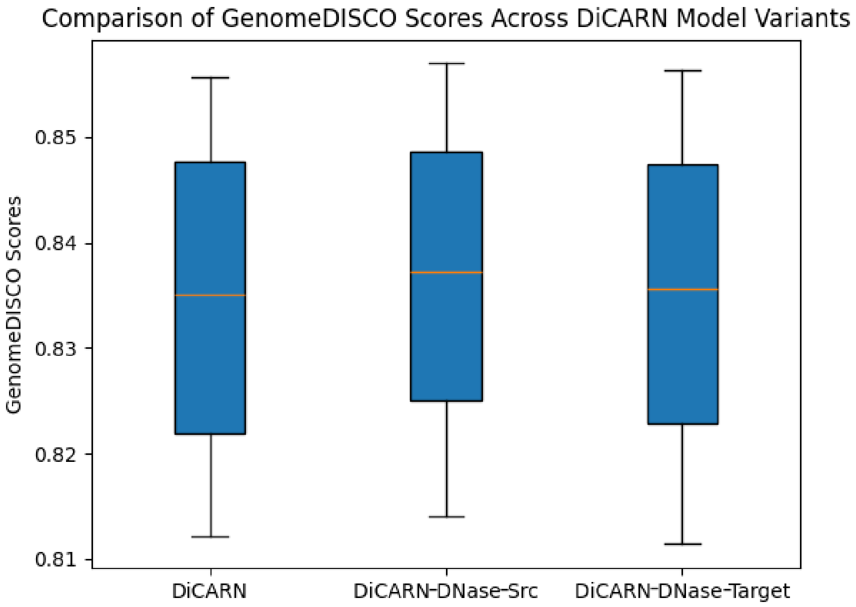


Figure 4: The biological concordance score using GenomeDISCO for DiCARN is contrasted against the Target DNase and Source DNase variants. These results are based on the 1/16 downsampled K562 dataset.

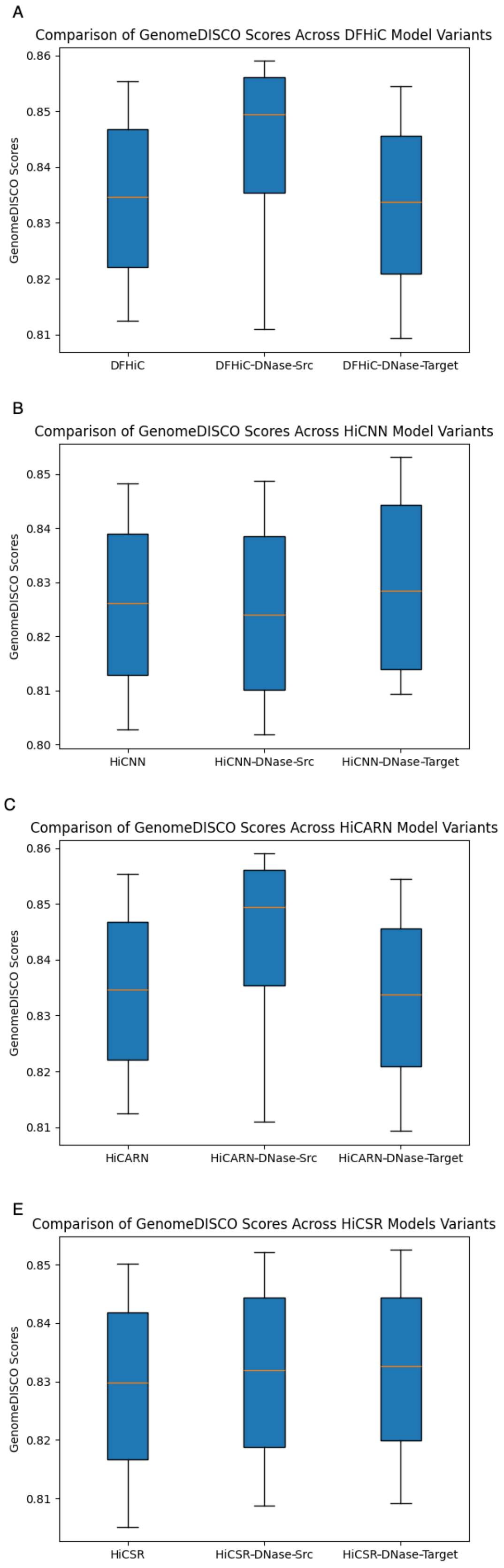


Figure 5: The biological concordance score using GenomeDISCO for all other vanilla models is compared to their corresponding DNase-induced version. This experiment is performed using the 1/16 downsampled K562 datasets.

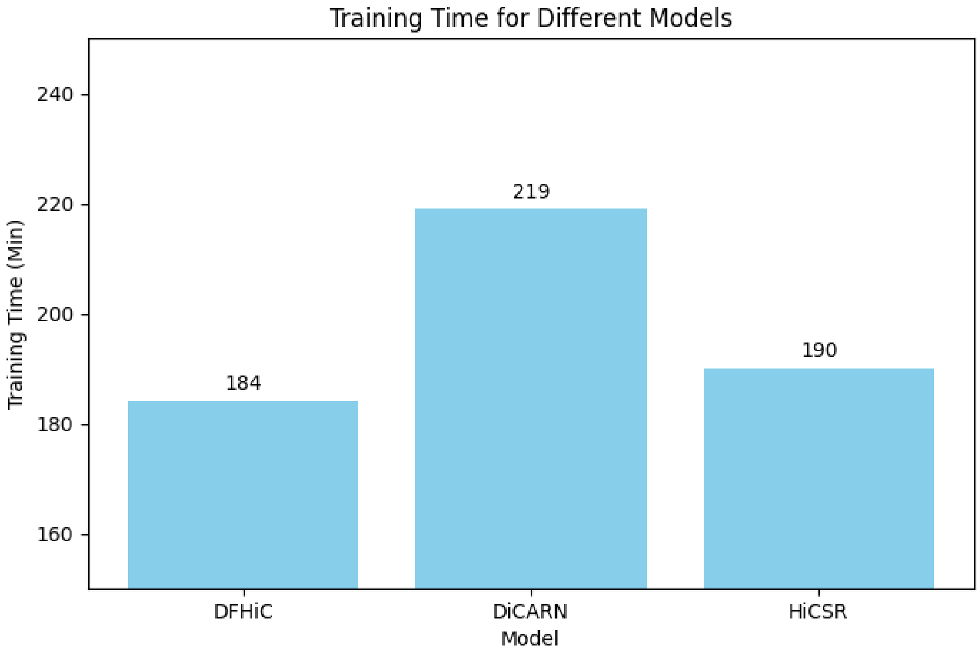


Figure 6: Time taken to train each model per 100 epochs.

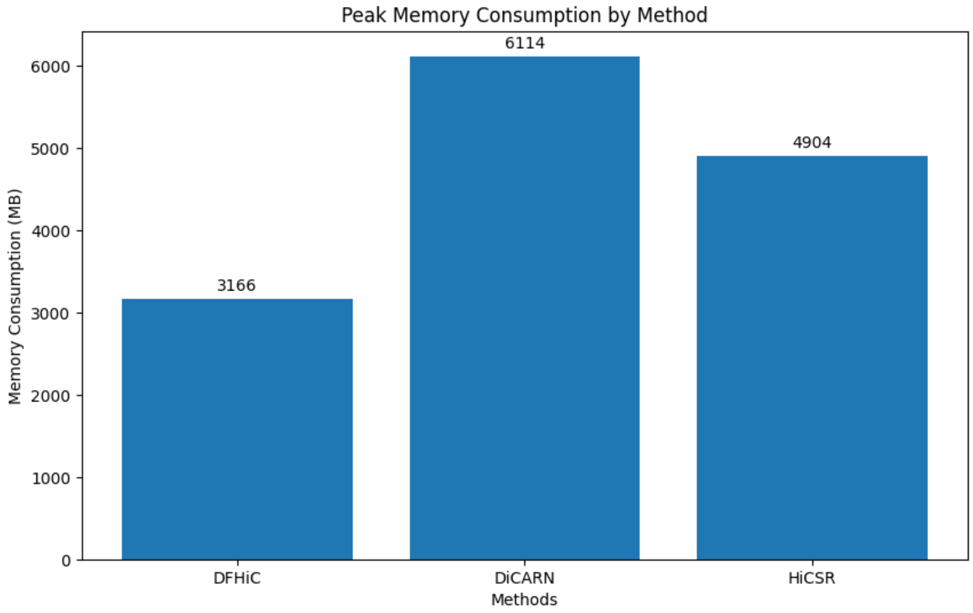


Figure 7: Peak memory consumption by each model at training.
